# Closely related *Vibrio alginolyticus* strains encode an identical repertoire of prophages and filamentous phages

**DOI:** 10.1101/859181

**Authors:** Cynthia Maria Chibani, Robert Hertel, Michael Hoppert, Heiko Liesegang, Carolin Charlotte Wendling

## Abstract

Filamentous vibriophages represent a massive repertoire of virulence factors which can be transferred across species boundaries, leading to the emergence of deadly pathogens. All filamentous vibriophages that were characterized until today were isolated from human pathogens. Considering frequent horizontal gene transfer among vibrios, we predict that other environmental isolates, including non-human pathogens also carry filamentous phages, of which some may encode virulence factors.

The aim of this study was to characterize the phage repertoire, consisting of prophages and filamentous phages, of a marine pathogen, *Vibrio alginolyticus*. To do so, we sequenced eight different *V. alginolyticus* strains, isolated from different pipefish and characterised their phage repertoire using a combination of morphological analyses and comparative genomics.

We were able to identify a total of five novel phage regions (three different *Caudovirales* and two different *Inoviridae*), whereby only those two loci predicted to correspond to filamentous phages (family *Inoviridae*) represent actively replicating phages. Unique for this study was that all eight host strains, which were isolated from different eukaryotic hosts have identical bacteriophages, suggesting a clonal expansion of this strain after the phages had been acquired by a common ancestor. We further found that co-occurrence of two different filamentous phages leads to within-host competition resulting in reduced phage replication by one of the two phages. One of the two filamentous phages encoded two virulence genes (Ace and Zot), homologous to those encoded on the *V. cholerae* phage CTXΦ. The coverage of these zot-encoding phages correlated positively with virulence (measured in controlled infection experiments on the eukaryotic host), suggesting that this phages is an important virulence determinant.

**Impact statement:** Many bacteria of the genus *Vibrio*, such as *V. cholerae* or *V. parahaemolyticus* impose a strong threat to human health. Often, small viruses, known as filamentous phages encode virulence genes. Upon infecting a bacterial cell, these phages can transform a previously harmless bacterium into a deadly pathogen. While filamentous phages and their virulence factors are well-characterized for human pathogenic vibrios, filamentous phages of marine vibrios, pathogenic for a wide range of marine organisms, are predicted to carry virulence factors, but have so far not been characterized in depth. Using whole genome sequencing and comparative genomics of phages isolated from a marine fish pathogen *V. alginolyticus*, we show that also environmental strains harbour filamentous phages that carry virulence genes. These phages were most likely acquired from other vibrios by a process known as horizontal gene transfer. We found that these phages are identical across eight different pathogenic *V. alginolyticus* strains, suggesting that they have been acquired by a common ancestor before a clonal expansion of this ecotype took place. The phages characterized in this study have not been described before and are unique for the Kiel *V. alginolyticus* ecotype.

**Data Summary:**

1. The GenBank accession numbers for all genomic sequence data analysed in the present study can be found in Table S1.
2. All phage regions identified by PHASTER analysis of each chromosome and the respective coverage of active phage loci are listed in Table S2.
3. GenBank files were deposited at NCBI for the two actively replicating filamentous phages VALGΦ6 (Accession number: MN719123) and VALGΦ8 (Accession number: MN690600)
4. The virulence data from the infection experiments have been deposited at PANGAEA: Accession number will be provided upon acceptance of the manuscript.

**Data statement:** All supporting data have been provided within the article or through supplementary data files. Four supplementary tables and six supplementary figures are available with the online version of this article.

## Introduction

Bacteriophages contribute significantly to bacterial adaptation and evolution. In particular through bacterial lysis and subsequent killing, phages impose a strong selection pressure on their bacterial hosts. However, phages can also transfer genetic material to neighbouring cells via horizontal gene transfer (HGT) thereby increasing bacterial genome plasticity (1). In addition, many phages, in particular temperate and filamentous phages, often carry virulence genes (2, 3). When such filamentous phages integrate into the bacterial chromosome, they can alter the phenotype of their bacterial host, resulting in increased bacterial virulence, through a process known as lysogenic conversion (3).

One of the best-known examples of lysogenic conversion is the transformation of non-toxigenic *Vibrio cholerae* into deadly pathogens via the filamentous CTXΦ phage, that carries the cholera toxin (CT) (3). Since this first description of lysogenic conversion in 1996, several other filamentous phages of which many carry bacterial virulence factors have been discovered in particular for the genus *Vibrio*. For instance, two phages VfO4K68 and VfO3K6 that carry the zona occludens toxin (Zot) and the accessory cholera enterotoxin (Ace), have been isolated from *V. parahaemolyticus* (4, 5). Zot and Ace are particularly common among vibriophages isolated from human pathogens, such as *V. cholerae* and *V. parahaemolyticus* (6-8) but are also present in prophage-like elements of non-human pathogens such as *V. coralliilyticus* (9) and *V. anguillarum* (10), suggesting frequent HGT among different vibrio species (11).

Other environmental vibrios, which cause severe diseases, not only in marine animals but also in humans, include for instance: *V. splendidus, V. tubiashii* and *V. alginolyticus* (12-15). While virulence of these vibrio species is often attributed to multiple factors, such as temperature and host immunity (16), the phage repertoire and any phage-encoded virulence factors of environmental isolates are often not well characterized. One reason might be that only long-read sequencing data which allow us to generate fully-closed genomes, are suitable to reliably identify integrated phages. Another reason might be a research bias towards human pathogens. There are 196 closed *Vibrio* genomes (as of October 2019), of which more than 50% comprise human pathogens such as *V. cholerae* (51 genomes), *V. parahaemolyticus* (33 genomes), and *V. vulnificus* (18 genomes), while all other environmental isolates are represented with fewer than 10 genomes per species. Additionally, to our knowledge only 17 filamentous vibriophages have so far been described in detail of which all were isolated from human pathogens, i.e. *V. cholerae* (11 phages) and *V. parahaemolyticus* (5 phages). Some of these 17 filamentous phages, encode at least one virulence factor. Other filamentous vibriophages, which do not encode virulence factor, are still able to transfer them to other strains by means of specialized transduction (17, 18). Recombination between filamentous vibriophages can further result in hybrid phages, which are then able to vertically transmit toxins to other vibrios (17, 19). Indeed, some filamentous vibriophages can infect distantly related species (20) and phage-mediated horizontal transfer of virulence genes seems to be a dynamic property among environmental vibrios (21). Thus, the low number of well-described filamentous vibriophages is concerning and it is essential that we start to characterize filamentous phages and their virulence factors in environmental *Vibrio* isolates. We predict, that also filamentous phages isolated from environmental vibrios, may contain virulence factors responsible for disease outbreaks in marine eukaryotes. Indeed, a closer look at more than 1,800 *Vibrio* genome sequences, covering 64 species revealed that 45% harboured filamentous phages which encoded Zot-like proteins (22). Even though a detailed characterization of these phages is missing, this study suggests that also filamentous vibriophages of non-human pathogens contain virulence genes in (22).

The aim of the present study was to identify prophages and filamentous phages (both types will also be referred to as non-lytic phages in the present study) isolated from closely related *V. alginolyticus* strains and to characterize their virulence potential. *V. alginolyticus*, a ubiquitous marine opportunistic pathogen can cause mass mortalities in shellfish, shrimp and fish, resulting in severe economic losses worldwide (23-25). Additionally, wound infections and fatal septicaemia in immunocompromised patients caused by *V. alginolyticus* have been reported in humans (26). In contrast to the classical human pathogenic vibrios, only little is known about *V. alginolyticus* phages and their potential role in its virulence. By combining morphological and comparative genomic analyses we identified and characterized three prophages and two novel filamentous phages from eight different environmental *V. alginolyticus* isolates of which one filamentous phage encoded virulence genes homologues to those found in *V. cholerae* and *V. parahaemolyticus*.

## Methods

*Vibrio alginolyticus* strains used in the present study were isolated either from the gut or the gills of six different pipefish (*Syngnathus typhle*) in the Kiel Fjord in 2012 (27) and have been shown to cause mortality in juvenile pipefish ((28), and this study). Using a combination of PacBio and Illumina sequencing we generated eight closed bacterial genomes of the host bacteria as described in (28).

### Phage isolation and sequencing

#### Prophage induction

We induced filamentous phages from all nine *V. alginolyticus* strains using Mitomycin C (Sigma), for details see (28), with some modifications: bacteria were grown in liquid Medium101 (Medium101: 0.5% (w/v) peptone, 0.3% (w/v) meat extract, 3.0% (w/v) NaCl in MilliQ water) at 250 rpm and 25 °C overnight. Cultures were diluted 1:100 in fresh medium at a total volume of 20ml and grown for another 2h at 250 rpm and 25 °C to bring cultures into exponential growth before adding Mitomycin C at a final concentration of 0.5 μg/ml. Afterwards, samples were incubated for 4 h at 25 °C at 230 rpm. After 4 h, lysates were centrifuged at 2500 *g* for 5 min. The supernatant was sterile filtered using 0.45 µm pore size filter (Sarstedt, Nümbrecht, Germany). We added lysozyme from chicken egg white (10µg/ml, SERVA Heidelberg, Germany) to disrupt the cell walls of potentially remaining host cells, RNAse A (Quiagen, Hilden, Germany) and DNAse I (Roche Diagnostics, Mannheim, Germany) at a final concentration of 10 µg/ml to remove free nucleic acids and remaining host cells as described in (29). After incubation at 25°C for 16 hours phage particles were sedimented by ultracentrifugation using a Sorvall Ultracentrifuge OTD50B with a 60Ti rotor applying 200,000 g for 4 hours. The supernatant was discarded and the pellet was dissolved in 200 µl TMK buffer, and directly used for DNA isolation.

#### Prophage DNA extraction

DNA isolation was performed using a MasterPure DNA Purification kit from Epicenter (Madison, WI, USA). We added 200 µl 2x T&C-Lysis solution containing 1 µl Proteinase K to the phage suspensions and centrifuged the samples for 10 min at 10,000 g. The supernatant was transferred to a new tube, mixed with 670 µl cold isopropanol and incubated for 10 min at – 20°C. DNA precipitation was performed by centrifugation for 10 min at 17,000 g and 4°C. The DNA pellet was washed twice with 150 µl 75% ethanol, air-dried and re-suspended in DNase free water.

#### Prophage sequencing

dsDNA for library construction was generated from viral ssDNA in a 50 µl reaction. The reaction was supplemented with 250 ng viral ss/DNA dissolved in water, 5 pmol random hexamer primer (#SO142, Thermo Scientific), 10 units Klenow Fragment (#EP0051, Thermo Scientific) and 5 µmol dNTPs each (#R0181, Thermo Scientific) and incubated at 37°C for 2 hours. The reaction was stopped by adding 1 µl of a 0.5M EDTA pH 8 solution. The generated DNA was precipitated by adding 5 µl of a 3M sodium acetate pH 5.2 and 50 µl 100% Isopropanol to the DNA solution, gently mixing and chilling for 20 min at −70°C. DNA was pelleted via centrifugation at 17,000 g, 4°C for 10 min. Pellets were washed twice with 70% ethanol. Remaining primers and viral ss/DNA were removed in a 50 µl reaction using 10 units S1 nuclease (#EN0321, Thermo Scientific) for 30 min at 25°C. S1 nuclease was inactivated through addition of 1µM 0.5M EDTA pH 8 and incubation for 10 min at 70°C. Consequently ds/DNA was precipitated as described above and resolved in pure water. Presence of ds/DNA was verified via TAE gel electrophoresis in combination with an ethidium bromide staining and visualization via UV-light. NGS libraries were generated with the Nextera XT DNA Sample Preparation Kit (Illumina, San Diego, USA), and the sequencing was performed on an Illumina GAII sequencer (Illumina, San Diego, USA). All generated reads were checked for quality using the programs FastQC (30) and Trimmomatic (31).

### Transmission electron

Electron microscopy was carried out on a Jeol 1011 electron microscope (Eching, Germany). Negative staining and transmission electron microscopy (TEM) of phage-containing particles was performed as described previously (29, 32). Phosphotungstic acid dissolved in pure water (3%; pH 7) served as staining solution.

### Genomic analysis

#### Prediction of phage regions

All host genomes were scanned with PHASTER (33) to identify prophage like elements in each chromosome. Predicted prophage regions were further analysed using Easyfig (34) for pairwise phage sequence comparisons and synteny comparisons with an *E*-value cut-off of 1e−10. We used SnapGene Viewer (v. 4.3.10) to generated circular genomes including the predicted prophage regions from each strain.

#### Annotation

Annotation was performed using Prokka v1.11 (35) which was applied using prodigal for gene calling (36), *Vibrio* as the genus reference (--genus *Vibrio* option) and a comprehensive *Inoviridae* vibriophage protein database as a phage features reference database. Reference *Inoviridae* vibriophages used for the reference protein fasta database are listed in (Supplementary material, Table S3).

#### Prediction of active phage regions

All reads from phage DNA have been mapped using bowtie2 (37) to the corresponding reference *V. alginolyticus* genome. The generated mapping files were analysed using TraV (38) to visualize phage DNA derived coverage within the genomic context. Increased coverage was exclusively observed in genomic regions that have been identified by PHASTER (33) as phage regions. This was used as an indication for active prophages.

To estimate the relative phage production of each active phage locus we estimated the coverage of each locus relative to the coverage of the chromosome. Deeptools v.3.3.0 (39) was used to compute read coverage which was normalized using the RPKM method as follows: RPKM (per bin) = number of reads per bin/ (number of mapped reads (in millions) * bin length (kb)). The length of the bin used is 1kb.

#### Comparative genomic analysis

We used the MUSCLE algorithm implemented in AliView v. 1.15 (40) to conduct whole genome alignments within all phage-groups that showed a high similarity based on Easyfig. Additionally, we performed alignments of the flanking regions by comparing five genes located upstream and five genes located downstream of each integrated phage.

To investigate the phylogenetic relationship of *Vibrio* phage VALGΦ6 and *Vibrio* phage VALGΦ8 with other well-studied filamentous phages we generated a phylogenetic tree based on the major coat protein (pVIII) of 20 well-characterized filamentous phages, which determined the structure of the virion coat. This protein is the most abundant protein present in all filamentous phages (41) and commonly used to infer phylogenetic relationships between filamentous phages. After alignment of the protein sequences using MUSCLE (42), we constructed a phylogenetic tree using the Bayesian Markov chain Monte Carlo (MCMC) method as implemented in MrBayes version 3.2.5 (43, 44). The TN93 (45) model plus invariant sites (TN93 + I), as suggested by the Akaike information criterion (AIC) given by jModelTest (46), was used as statistical model for nucleotide substitution. The MCMC process was repeated for 10^6^ generations and sampled every 5000 generations. The first 2000 trees were deleted as burn-in processes and the consensus tree was constructed from the remaining trees. Convergence was assured via the standard deviation of split frequencies (<0.01) and the potential scale reduction factor (PSRF∼1). The resulting phylogenetic tree and associated posterior probabilities were illustrated using FigTree version 1.4.2 (http://tree.bio.ed.ac.uk/software/figtree/). We used *Propionibacterium phage B5* which is a phage preying on a gram-positive bacterium as an outgroup.

We additionally compared the predicted phage regions from the present study with potential phage regions from all other so far published fully closed *V. alginolyticus* genomes (Table S4). To do so, we identified potential phage regions on each chromosome using PHASTER (33) and compared those with the phage regions from the present study using Easyfig (34).

#### Analysis of virulence factors

We found that one of the active filamentous phages (i.e. *Vibrio* phage VALGΦ6) contains the virulence cassette comprising the Zot and the Ace proteins, which is frequently found in vibriophages and responsible for severe gastro-intestinal diseases (47, 48). To compare these two proteins with other Zot and Ace proteins isolated from various vibriophages we generated protein alignments using AliView (40) and examined the presence of Walker A and Walker B motifs in *Vibrio* phage VALGΦ6 Zot proteins. We further used the TMHMM Server (http://www.cbs.dtu.dk/services/TMHMM/) to confirm the presence of a transmembrane domain typically found in the Zot protein.

#### Infection experiments

We performed a controlled infection experiment to estimate the virulence of the eight sequenced strains on juvenile pipefish (a detailed description of methods and the statistical analysis can be found in (28). Briefly, we fed 9-12 juvenile pipefish per tank in using triplicate tanks with *Artemia* nauplii which were previously exposed to ∼10^9^ CFU/ml or seawater as control. Twenty-four hours post infection, each fish was killed and bacterial load was determined as colony forming units (CFU/ml) as in (28).

GenBank files were deposited at NCBI for the two actively replicating filamentous phages VALGΦ6 (Accession number: MN719123) and VALGΦ8 (Accession number: MN690600)

## Results

### 1. General overview

We sequenced the bacterial DNA of eight closely related *Vibrio alginolyticus* strains as well as the DNA extracted from the supernatant of mitomycin C treated liquid cultures of each strain. Within the eight sequenced *V. alginolyticus* strains we discovered three different prophage regions each of which could be assigned to the family *Caudovirales* and two different regions that were assigned to filamentous phages (family *Inoviridae*). From the sequenced supernatant we could only identify filamentous phages but no head-tail phages, suggesting that in the present strains, filamentous phages are the only active replicating phages. To locate the exact positions of the induced prophages, we performed a PHAGE-seq experiment (29). In control experiments, the complete procedure has been applied without mitomycin C where the reference genomes were sequenced using Illumina technology. Both experiments revealed an increased coverage exclusively at *Inoviridae* loci (Supplementary material Figure S1). This indicates that induced and non-induced cultures produce comparable amounts of particles encoded by the same filamentous phage. As a further control total DNA without DNase A treatment resulted in a coverage increased by the factor of 100-100,000 at the loci encoding filamentous phages compared to the average chromosomal coverage. We thus conclude that the cultures produced a permanent amount of phage particle protected ssDNA independent of the induction from mitomycin C.

### 2. Caudovirales

Whole genome comparison between the eight sequenced *V. alginolyticus* strains revealed the presence of three different prophage regions belonging to the family *Caudovirales*, none of which generated phage particles nor protein protected DNA in the experimental settings used in this study (Figure 1). Thus, a more thorough classification based on morphological characterization was not possible. We further did not find regions of increased coverage for these three *Caudovirales* regions (Supplementary material Figure S1) on the bacterial chromosomes indicating that these phages were neither actively replicating in uninduced bacterial cultures nor able to switch to the lytic cycle upon induction with mitomycin C. We could not identify sequence similarities between these three different *Caudovirales* phages, suggesting that they are genetically distinct phages. However, each of the three *Caudovirales* phages was 100% identical across all eight strains where they all have the same integration site (Figure 2).

**Figure 1.**
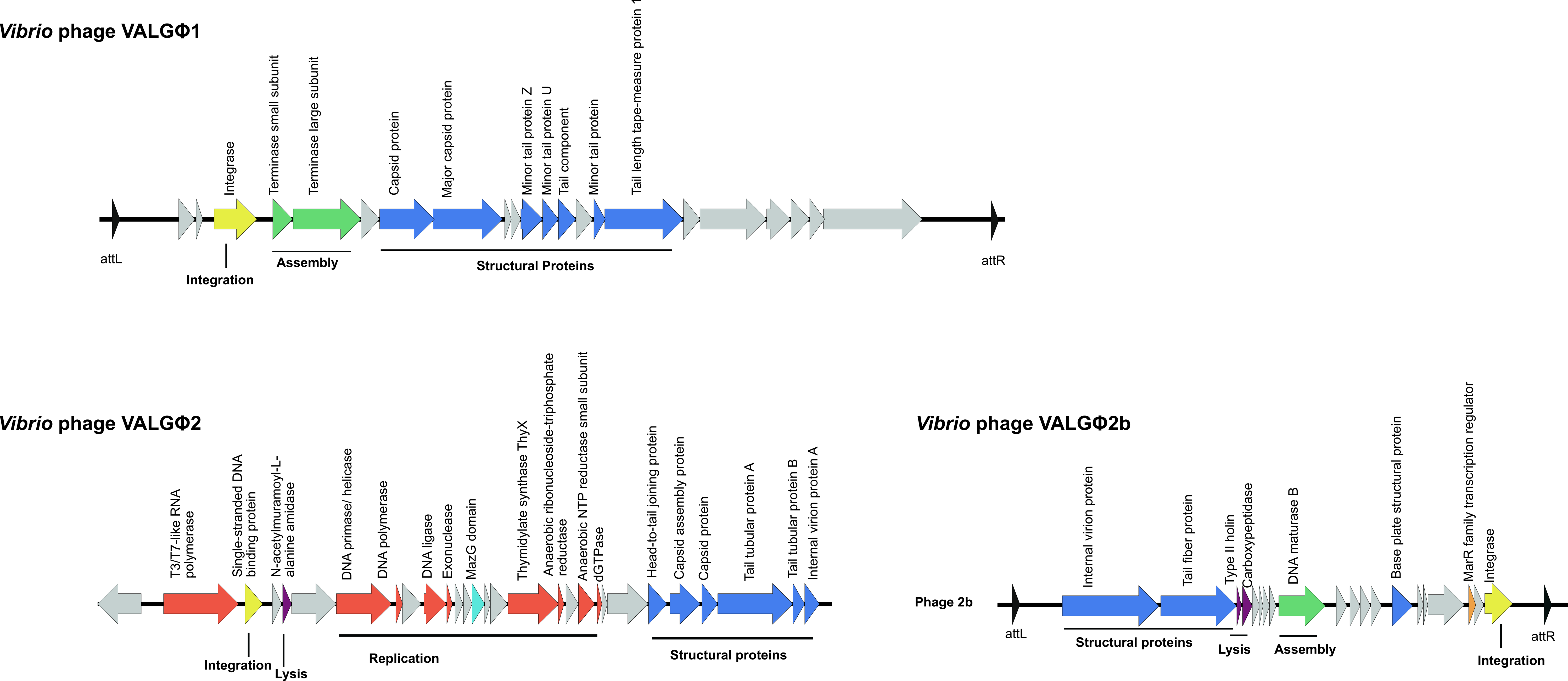
Genomic maps of *Vibrio* phage VALGΦ1 (top), *Vibrio* phage VALGΦ2 (bottom left) and *Vibrio* phage VALGΦ2b (bottom right). ORFs are color-coded according to predicted function: red: replication, green: assembly, blue: structural proteins, yellow: integration, purple: lysis, orange: accessory genes, grey: hypothetical proteins.

**Figure 2.**
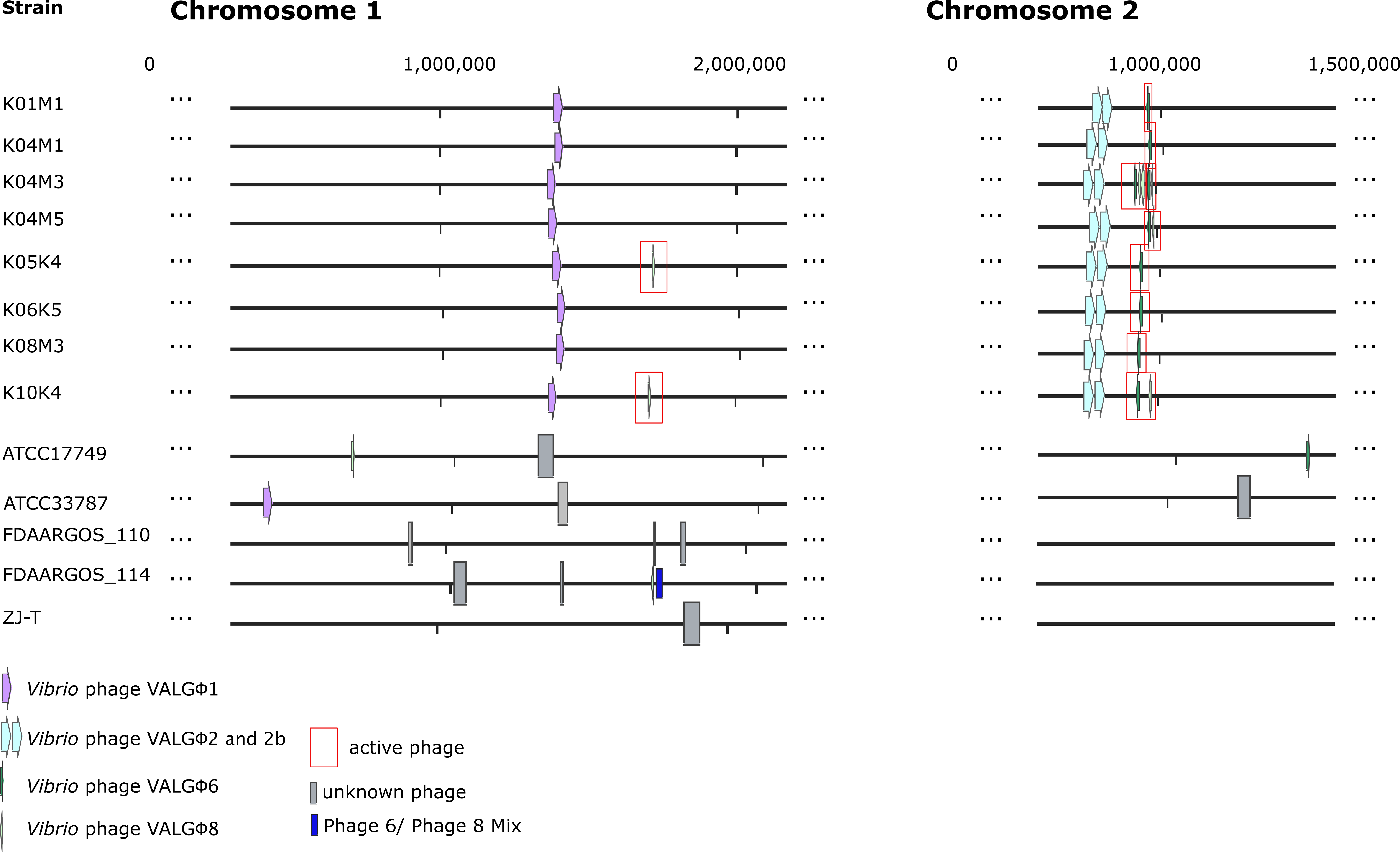
Whole Chromosome alignment with prophage regions in coloured boxes/ arrows. Active prophages are marked in red. Blocks of the same colour indicate phage-types, purple: *Vibrio* phage VALGΦ1, light blue: *Vibrio* phage VALGΦ2, *Vibrio* phage VALGΦ2b complex, dark green: *Vibrio* phage VALGΦ6, light green: *Vibrio* phage VALGΦ8, dark blue: multi-phage complex containing sequences of *Vibrio* phage VALGΦ6 and *Vibrio* phage VALGΦ8, grey: unknown phages.

#### *Vibrio* phage VALGΦ1

The genome of *Vibrio* phage VALGΦ1 is composed of a 33.3kb DNA molecule with a GC content of 46.06 % and no tRNAs. The total open reading frames (ORFs) is 22, with 10 ORFs assigned to one of five functional groups typical for phages (Replication, Assembly, Structural proteins, Integration, Lysis) and 12 ORFs to hypothetical proteins (Figure 1). All ORFs were orientated in the same direction. Even though *Vibrio* phage VALGΦ1 could not be found in induced and uninduced supernatants, it is predicted to be intact according to PHASTER. *Vibrio* phage VALGΦ1 is exclusively found on Chromosome 1, where it has a unique integration site, which is identical across all eight sequenced strains. The phage genome as well as the flanking regions (five genes upstream and five genes downstream of the integrated phage) showed 100% sequence similarity across all eight sequenced strains, suggesting that the phage is highly conserved across host-strains. Comparative genomic analysis between *Vibrio* phage VALGΦ1 and ten closest hits on NCBI reveals that the two closest related phages are FDAARGOS_105 integrated on chromosome 1 of *V. diabolicus* with a query cover of 77% and a similarity of 94.68% followed by an uncharacterized region on chromosome 1 of *V. alginolyticus* ATCC 33787 with a query cover of 57% and a similarity of 96.13%. These low query covers suggest that *Vibrio* phage VALGΦ1 is a novel bacteriophage.

#### *Vibrio* phage VALGΦ2

The genome of *Vibrio* phage VALGΦ2 is 26.3 kb with a GC content of 49.37 %, no tRNAs and a total of 29 ORFs, with 22 assigned to one of five functional phage-related groups and seven hypothetical proteins (Figure 1). All ORFs were orientated in the same direction. *Vibrio* phage VALGΦ2 is predicted to be questionable by PHASTER, suggesting that it does not contain sufficient prophage genes to be considered a complete functional phage. Even though, *Vibrio* phage VALGΦ2 has a unique integration site on chromosome 2 across all eight strains, the upstream region is not identical across strains. In contrast, the downstream region of *Vibrio* phage VALGΦ2 is identical across all strains and has a length of 2584 bp followed by another prophage, identical across all sequenced strains and, identified as *Vibrio* phage VALGΦ2b. Due to their incompleteness and the short gap between these two phages we refer to them as a *Caudovirales* complex consisting of *Vibrio* phage VALGΦ2 and *Vibrio* phage VALGΦ2b.

#### *Vibrio* phage VALGΦ2b

*Vibrio* phage VALGΦ2b is predicted to be incomplete by PHASTER, suggesting that it may represent a cryptic phage. The genome of *Vibrio* phage VALGΦ2 is 26.5 kb long, with zero tRNAs and a GC content of 48.32%. Of the 20 identified OFRs, 12 could be assigned to phage-functional groups and eight as hypothetical proteins. *Vibrio* phage VALGΦ2 contains an ORF assigned as MarR family transcriptional regulator accompanied with a transposase 3749 bp upstream.

### 3. Inoviridae

#### Phage morphology

We determined the morphology of all active phages from every strain using a transmission electron microscope (TEM, see Supplementary material, Figure S2). According to the International Committee on Taxonomy of Viruses (ICTV), all phage particles were identified as filamentous phages.

#### Phage genomics

Within the eight sequenced *V. alginolyticus* strains we could identify two different filamentous phages, i.e. *Vibrio* phage VALGΦ6 and *Vibrio* phage VALGΦ8. Both phages contain single-stranded ssDNA genomes of 8.5 and 7.3 kbp in size and a GC content of 44.6% and 46.3%, respectively. ORFs were mostly orientated in a single direction, whereas the transcription regulator was transcribed in the reverse direction (Figure 3). Both phages showed similar functional genes (typical for *Inoviridae*), which could be roughly grouped into three functional modules: Replication, assembly or structural proteins (41). *Vibrio* phage VALGΦ6 and *Vibrio* phage VALGΦ8 share relatively little homology, except for proteins involved in DNA replication (Figure 3).

**Figure 3.**
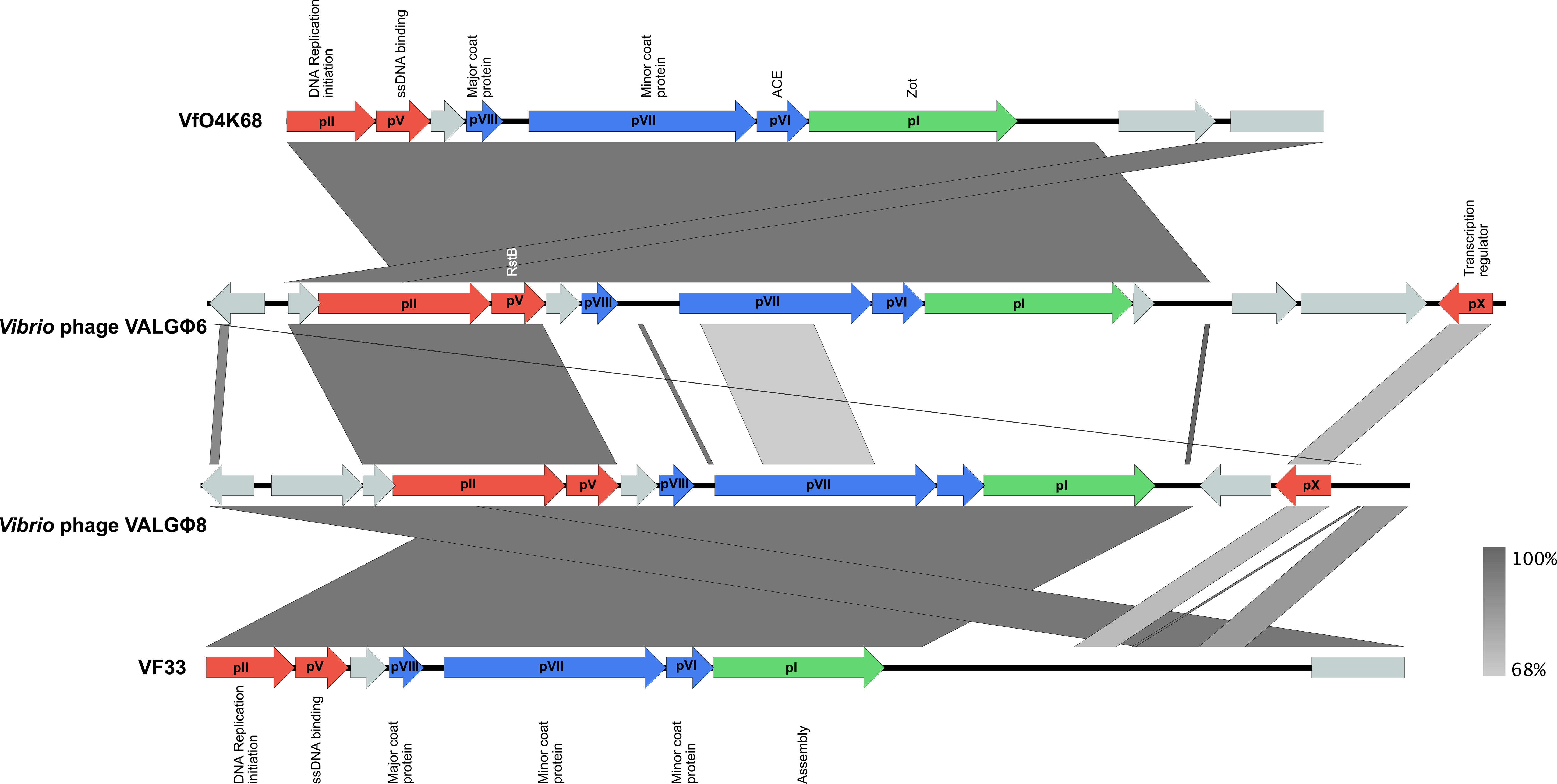
Genomic maps of *Vibrio* phage VALGΦ6 (second from top) and *Vibrio* phage VALGΦ8 (third from top) in comparison to VfO4K68 (top) and VF33 (bottom). ORFs are color-coded according to predicted function: red: replication, green: assembly, blue: structural proteins, grey: hypothetical proteins. High homologous sequences are indicated by dark grey and low homologous sequences by light grey. pI – pX correspond to known filamentous phage proteins and putative homologues.

*Vibrio* phage VALGΦ6 can be found exclusively on chromosome 2 in all eight strains, has a unique integration site and is identical across all strains. In contrast, *Vibrio* phage VALGΦ8 can only be found in five out of the eight strains and has a more diverse life-style. It can integrate on chromosome 2 (strain K04M3, K04M5, K10K4), chromosome 1 (Strain K05K4, K10K4) or exists extra-chromosomally (strain K04M1 without an intrachromosomal copy or in strain K05K4 with an intrachromosomal copy, Figure 4). When integrated on chromosome 2, *Vibrio* phage VALGΦ8 is always located directly behind *Vibrio* phage VALGΦ6, sometimes resulting in multi-phage cassettes (Figure 2). When integrated on chromosome 1, *Vibrio* phage VALGΦ8 is orientated in a reverse order as on chromosome 2. In strain K10K4, *Vibrio* phage VALGΦ8 is found on both chromosomes.

**Figure 4.**
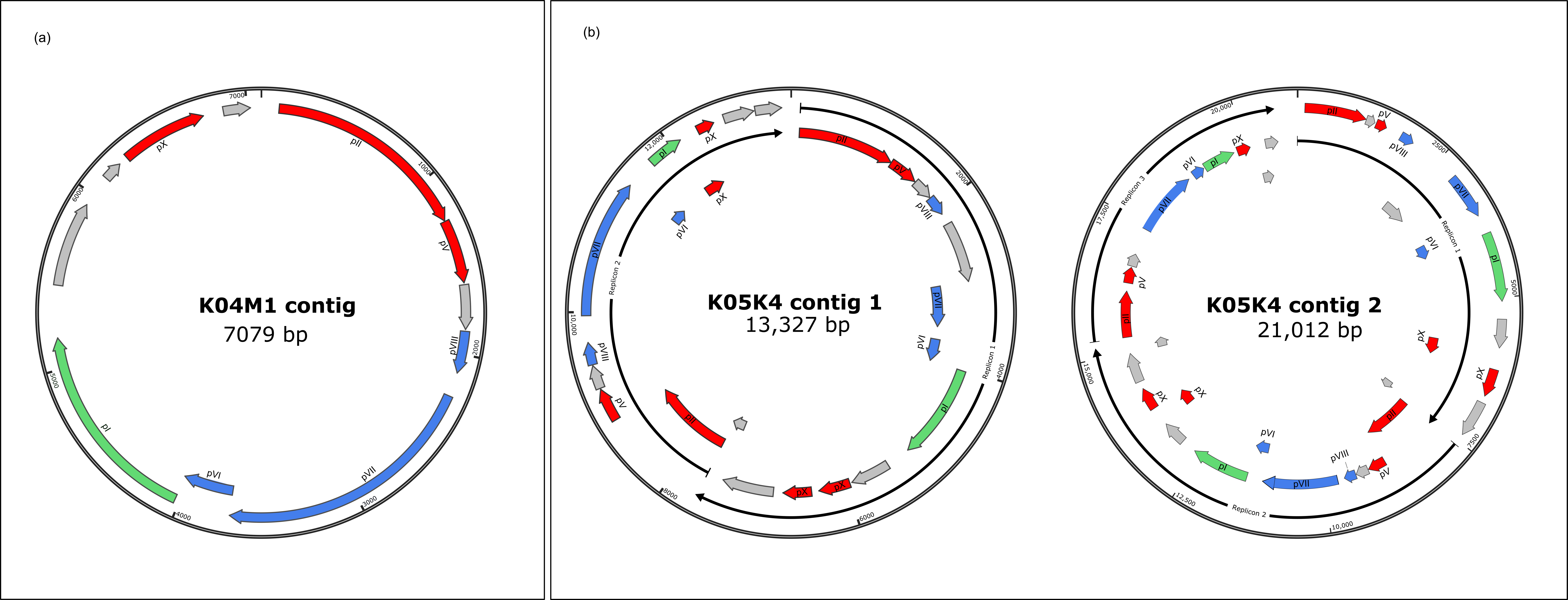
Extrachromosomal contigs of (a) Strain K04M1 and (b) Strain K05K4 with the 2-replicon containing contig (left) and the 3-replicon containing contig (right). ORF-coding and protein names as in Figure 3.

#### Phage activity

All loci predicted to correspond to filamentous phages represent actively replicating phages. We conclude this from several lines of evidence. First, we were able to detect filamentous phages in TEM pictures of all cultures (see Supplementary material, Figure S2). Second, phage particles isolated from induced and uninduced cultures contained exclusively DNA that matched at these phage loci (Supplementary material, Figure S1, Table S2). Differences in coverage values of loci corresponding to both filamentous phages we found that the production of phage particles varies across phage regions and strains (Supplementary material, Table S2). For strains that did not contain *Vibrio* phage VALGΦ8, we found that all regions encoding *Vibrio* phage VALGΦ6 had on average a 100000x higher coverage relative to the coverage of the chromosome. However, the presence of *Vibrio* phage VALGΦ8 reduced the coverage of *Vibrio* phage Vibrio phage VALGΦ6 encoding regions by 10 – 1000x, but only when *Vibrio* phage VALGΦ8 was integrated on chromosome 2, not when it existed exclusively extrachromosomal or had an additional copy on chromosome 1.

#### Phylogeny

Whole genome alignment (Figure 3) and phylogenetic comparisons (Figure 5) based on the major coat protein (pVIII) suggest that *Vibrio* phage VALGΦ6 and *Vibrio* phage VALGΦ8 group closely with other known filamentous vibriophages. Overall, filamentous vibriophages group more closely with class II phages of *Pseudomonas* and *Xanthomonas* and form a distinct cluster from class I filamentous coliphages. *Vibrio* phage VALGΦ6 shares more sequence homology with VfO4K68 and VfO3K6, both isolated from *V. parahaemolyticus* (Figure 3). *Vibrio* phage VALGΦ8 shares more sequence homology with VF33 also isolated from *V. parahaemolyticus*. Blastn comparisons using whole phage genomes suggest that *Vibrio* phage VALGΦ6 and *Vibrio* phage VALGΦ8 are different from other bacteriophages described until today. For *Vibrio* phage VALGΦ6 the two closest hits were the two *V. parahaemolyticus* phages VfO4k68 and VfO3k6 with query covers of 66% and 75% and similarity values of 94.65% for each phage. The closest hits for *Vibrio* phage VALGΦ8 were the two *V. parahaemolyticus* phages Vf12 and Vf13 with a query cover of 88% and a similarity of 94.65%.

**Figure 5.**
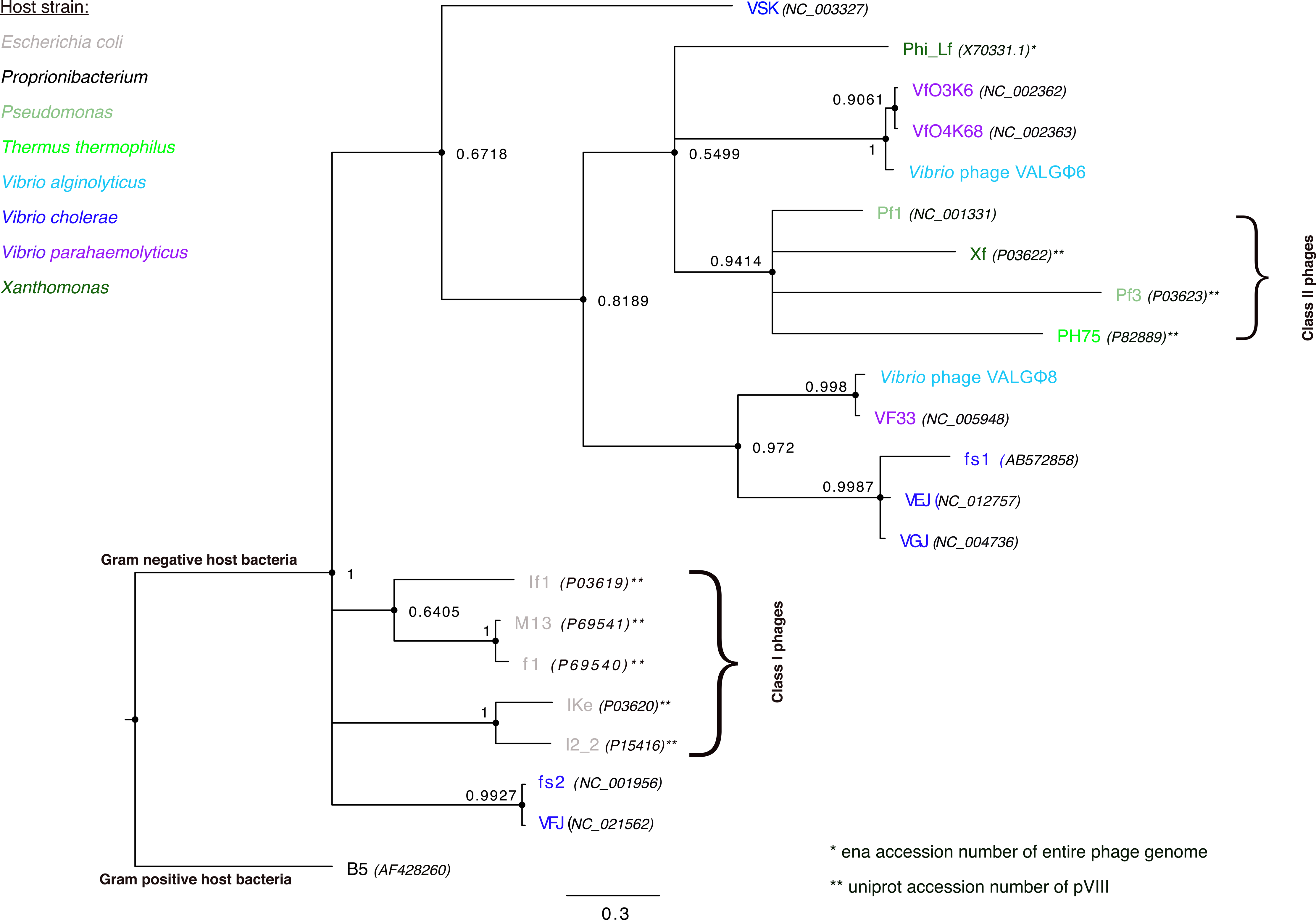
Phylogenetic tree based on the major coat protein (pVIII) alignment highlighting the position of *Vibrio* phage VALGΦ6 and *Vibrio* phage VALGΦ8 relative to other filamentous phages. The corresponding NCBI accession numbers for the different phages are denoted in brackets, ena accession numbers are indicated with an *, uniport accession numbers of the major coat protein are indicated with **. For outgroup *Proprionibacterium phage B5* was used. Class I and Class II phages represent clusters according to (62).

To compare all non-lytic phages between strains from the present study and other *V. alginolyticus* isolates we used PHASTER to predict prophages from all available closed non-Kiel *V. alginolyticus* genomes and found a total of 14 predicted prophage regions (Table S4). Comparisons between those uncharacterized vibriophages and phages from the present study revealed that *Vibrio* phage VALGΦ6 and the *Caudovirales* cassette consisting of *Vibrio* phage VALGΦ2 and 2b is unique to the Kiel *alginolyticus* system. In contrast, we found integrated *Inoviridae* with high similarity to *Vibrio* phage VALGΦ8 in four of the six non-Kiel *V. alginolyticus* strains and one integrated *Caudovirales* that shared high similarity with *Vibrio* phage VALGΦ1 and had the same attL/ attR sequence, i.e. CGTTATTGGCTAAGT (Figure 2). Despite having two, respectively one, unique integration site within the Kiel isolates, phages from the non-Kiel isolates with high similarity to *Vibrio* phage VALGΦ8 and *Vibrio* phage VALGΦ1 were mostly integrated on different positions in the respective chromosomes (Figure 2). All other uncharacterized phages did not contain functional genes typical for *Inoviridae* suggesting that no other filamentous phage is present in the non-Kiel strains. In contrast to the Kiel strains, where most phages were integrated on chromosome 2, only two out of the non-Kiel strains had prophages on chromosome 2 and one strain FDAARGOS_108 did not have a single prophage in its genome. Overall, the typical phage composition consisting of *Vibrio* phage VALGΦ6, *Vibrio* phage VALGΦ2 and 2b on chromosome 2 together with *Vibrio* phage VALGΦ1 on chromosome 1 is unique for our system and has not been found elsewhere.

### Multi-phage-cassettes

We found multi-phage cassettes on chromosome 2 in two strains, i.e. K04M3 and K04M5. While the cassette in K04M5 consists of *Vibrio* phage VALGΦ6 followed by *Vibrio* phage VALGΦ8, K04M3 has two multi-phage cassettes. The first cassette consists of *Vibrio* phage VALGΦ6, followed by a tandem repeat of two identical *Vibrio* phage VALGΦ8 regions, the second cassette, which is located 10389 bp downstream of the first cassette is identical to the one identified in K04M5. Even though *Vibrio* phage VALGΦ6 is identical across all strains, the second replicate in K04M3, which represents the start of the second multi-phage cassette, misses the transcription regulator and has major deletions, particularly affecting assembly and structural proteins (Supplementary material, Figure S4).

### Virulence of Kiel *V. alginolyticus* ecotypes

Comparative genomic analysis between virulence factors commonly encoded on filamentous phages revealed that only *Vibrio* phage VALGΦ6 contains the virulence cluster containing Ace and Zot. In contrast no known virulence factors could be found on *Vibrio* phage VALGΦ8 and the described *Caudovirales*. Sequence comparisons of Zot proteins encoded on different vibrios revealed that the *Vibrio* phage VALGΦ6 encoded Zot is highly similar to Zot genes encoded on other closely related *Vibrio* species from the *harveyi* clade (such as *V. parahaemolyticus* or *V. campbellii,* Figure S5). Even though we found less similarity between the *Vibrio* phage VALGΦ6 encoded Zot protein and CTXΦ-encoded Zot proteins, we found two conserved motifs (Walker A and B, common among human pathogens), which were at the N-terminal side of the Zot proteins (Figure S5). In addition, we found a transmembrane domain in the *Vibrio* phage VALGΦ6 encoded Zot protein (Figure S6), suggesting that similar to the CTXΦ-encoded Zot, the *Vibrio* phage VALGΦ6 encoded Zot is also a transmembrane protein.

Controlled infection experiments on juvenile pipefish revealed differences in total bacterial load among strains, a proxy for virulence. We found that those strains, that only encode *Vibrio* phage VALGΦ6 (i.e. K01M1, K06K5 and K08M3) were the most virulent (Figure 6). Whereas strains with a reduced coverage of *Vibrio* phage VALGΦ6 encoding regions were by far the least virulent strains, in particular strain K04M5, where we also observed the strongest reduction in coverage compared to *Vibrio* phage VALGΦ8-free strains.

**Figure 6.**
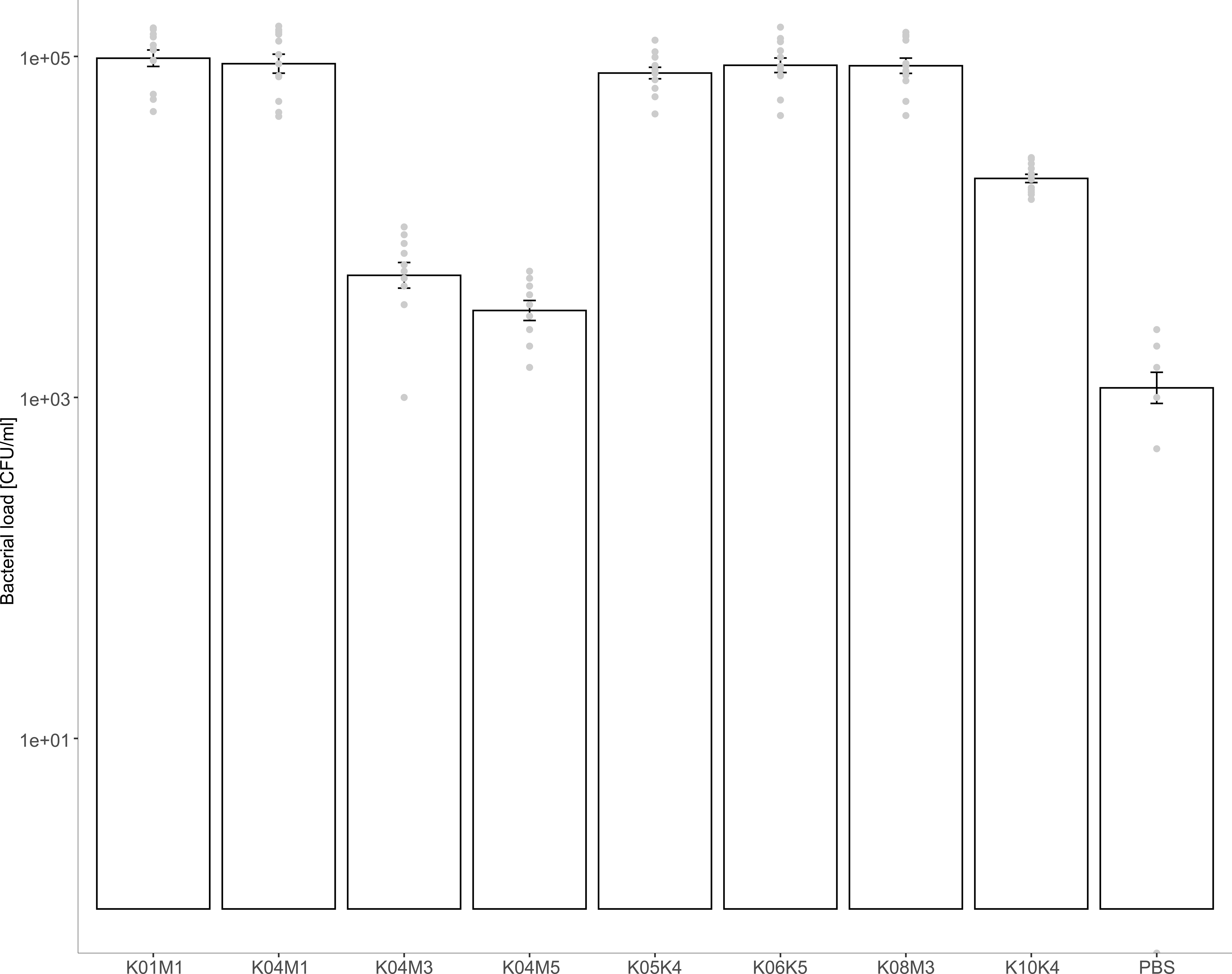
Virulence of all eight sequenced strains (x-axis), measured as bacterial load (CFU/ml).

## Discussion

We present five new non-lytic phages (comprising filamentous phages and prophages) isolated from eight different *Vibrio alginolyticus* strains. Using a combination of whole genome sequencing, comparative genomic analyses, and transmission electron microscopy we found three distinct phage regions belonging to the family *Caudovirales* and two distinct regions corresponding to actively replicating filamentous phages. Based on comparative genomic analyses we conclude that all five phages described in the present study are novel bacteriophages. Our main findings are that (1) closely related *V. alginolyticus* isolates, which were isolated from different eukaryotic hosts have identical bacteriophages, which are unique for this ecotype, (2) filamentous phages can have different life-styles and are able to supress each other, and (3) horizontal gene transfer (HGT) of *Vibrio* phage VALGΦ6 containing the virulence cluster comprising zona occludens toxin (Zot) and accessory cholera enterotoxin (Ace) may have led to the emergence of pathogenicity of the Kiel *V. alginolyticus* ecotype.

*Closely related V. alginolyticus isolates, which were isolated from different eukaryotic hosts share unique but identical bacteriophages*

Four of the five described phages in this study (i.e. *Vibrio* phage VALGΦ1 on chromosome 1, the *Caudovirales* complex consisting of *Vibrio* phage VALGΦ2 and VALGΦ2b as well as the filamentous *Vibrio* phage VALGΦ6 of chromosome 2) were present in all eight sequenced strains, had the same integration site and no variation in flanking regions on the chromosome (only exception: upstream region of *Vibrio* phage VALGΦ2). Only the filamentous *Vibrio* phage VALGΦ8 was not present in all strains, existed in two different life-styles (intra- and extrachromosomal) and had different integration sites (possible recombination with both chromosomes). All eight strains have no core genomic variation and sequence variation is mainly attributable to differences in mobile genetic elements (MGEs), such as plasmids and presence/ absence of *Vibrio* phage VALGΦ8 (49). Comparative genomic analyses across a wider range of *V. alginolyticus* isolates indicated that the phage repertoire of the Kiel *alginolyticus* ecotype is unique and cannot be found elsewhere. Thus, we hypothesize that the identical prophage composition in this ecotype together with the identical integration sites and flanking regions suggests that these phages may have been acquired from a common ancestor before a clonal expansion of the Kiel *alginolyticus* ecotype took place. Under this scenario we predict that these five prophages are increasing the fitness of this ecotype in the present habitat and are thus maintained by selection.

The sampling design, spanning two different organs (gills or gut) from six different pipefish allows us not only to look at the phage composition of closely related bacteria across eukaryotic hosts but also within eukaryotic hosts. We found more similarity within pipefish Nr. 4 (strains K04M1, K04M3 and K04M5) than across all six pipefish: First, all three strains contained *Vibrio* phage VALGΦ8, and and second, the only two multi-phage cassettes were found in strains K04M3 and K04M5, both isolated from pipefish Nr. 4. It is tempting to speculate that the high prevalence of *Vibrio* phage VALGΦ8 relative to all eight sequenced strains is a result of the close proximity between strains inside the gut, which favours the rapid horizontal spread of *Vibrio* phage VALGΦ8. Future experiments would be needed to study the likelihood for *Vibrio* phage VALGΦ8 to establish successful chronic infections and the circumstances which favour the different life-styles (extra- or intra-chromosomal) and integration sites (chromosome 1 or chromosome 2).

### Filamentous phages differ in their life-style

While *Vibrio* phage VALGΦ6 was exclusively found at one integration site across all eight sequenced strains (exceptions: the multi-phage cassettes in strains K04M3 and K04M5), *Vibrio* phage VALGΦ8 had different integration sites on both chromosomes and existed intra-and extrachromosomal. We identified one, respectively two extrachromosomal closed circular contigs within the assembly of strains K04M1 and K05K4 representing multimers of *Vibrio* phage VALGΦ8 (Figure 4). This indicates the presence of extrachromosomal phage replicons in two out of the eight sequenced *V. alginolyticus* genomes. Filamentous phages typically multiply via the rolling circle replication (RCR) mechanism (41). Considering that K05K4 contains another copy of *Vibrio* phage VALGΦ8 integrated on chromosome 1 and that the extrachromosomal contigs contain two, respectively three copies of *Vibrio* phage VALGΦ8 (Figure 4), we hypothesise that the K05K4 extrachromosomal contigs represent RCR intermediates of the integrated *Vibrio* phage VALGΦ8. However, to confirm or falsify this hypothesis experiments using knock-out versions of the intrachromosomal copy of *Vibrio* phage VALGΦ8 in strain K05K4 have to be performed which are beyond the scope of this study. In contrast, K04M1 does not contain an intrachromosomal version of *Vibrio* phage VALGΦ8 and the extrachromosomal contig of K04M1 only consists of one phage replicon (Figure 4). This suggests that *Vibrio* phage VALGΦ8 is able to establish a chronic extra-chromosomal infection without the need of an intrachromosomal copy.

### Within-host competition can lead to the reduction of phage producing particles

*Vibrio* phage VALGΦ1 has been predicted to be complete, but we did not find phage particles of head-tail phages in the supernatant nor did we detect any DNA-sequences in phage particles that map to its region in the chromosome under lab-conditions. Without a proof-of principle which would again require knock-out versions of the filamentous phages, we can only speculate that the *Caudovirales Vibrio* phage VALGΦ1 is supressed by the two actively replicating filamentous phages. Indeed, within-host competition between different *Caudovirales* has been found in other systems, for instance in *Bacillus licheniformis* (29). Alternatively, *Vibrio* phage VALGΦ1 might be not induced within the conditions of our experimental set up or could have been wrongly predicted to be complete by the software but is, however not able to actively replicate suggesting prophage domestication, which has also been predicted for *Vibrio* phage VALGΦ2 and 2b. When head-tail phages switch from the lysogenic to the lytic cycle they always kill their host. Selection for a strict repression of the lytic life cycle of prophage inactivation should thus be strong (50). Indeed, bacterial genomes have numerous defective prophages and prophage-derived elements (51, 52), which presumably originate from pervasive prophage domestication (50). By domesticating prophages, bacteria can evade the risk of getting lysed but are still able to maintain beneficial accessory genes, in the present case for instance the Mar family proteins encoded on the defective *Vibrio* phage VALGΦ2b, which encode transcriptional regulators involved in the expression of virulence, stress response and multi-drug resistance (53, 54). Another case of within-host competition between phages is the 10-x reduced coverage of *Vibrio* phage VALGΦ6 in strains where both filamentous phages were present. Again, we hypothesise, that one phage, in this case *Vibrio* phage VALGΦ8, negatively affects the replication of another phage, here *Vibrio* phage VALGΦ6. As above, to verify or falsify this hypothesis, knock-out versions of strains containing both filamentous phages would be required, as have been used in (29). Within-host competition is common among different head-tail prophages within the same host leading to strong selection for short lysis time (55). However, to the best of our knowledge, nothing is known about within-host competition among filamentous phages and whether filamentous phages are able to supress each other’s replication. However, studies on the classical biotype *V. cholerae* where CTXΦ was present as an array of two truncated, fused prophages found, that even though the cholera toxin in expressed, no viral particles are produced (56). Deficiencies in the array-structure and not mutations affecting individual CTXΦ genes have been suggested to be responsible for the absence of phage particle production. Similarly, we found that the strongest reduction in coverage for *Vibrio* phage VALGΦ6 encoding regions, where *Vibrio* phage VALGΦ6 was present as part of a multi-phage cassette, containing arrays of two or more adjacent filamentous phages (strain K04M3 and strain K04M5). As we also did not find any genomic differences among the different regions encoding for *Vibrio* phage VALGΦ6, it is tempting to speculate that similar deficiencies in array structure are causing the coverage reduction of *Vibrio* phage VALGΦ6 encoding regions. Alternatively, suppression of one phage by another could be the result of methylation leading to a less efficient or even inactivate phage particle production. Future studies unravelling within-host interactions of filamentous phages should elucidate whether such within-host competitions can influence the dynamics and evolutionary trajectories of filamentous phages.

### HGT of Vibrio phage VALGΦ6 containing the virulence cluster comprising Zot and Ace may *have led to the emergence of the pathogenic Kiel V. alginolyticus ecotype*

Filamentous phages are the most recognized vibriophages and present in almost every *Vibrio* genome sequenced to date (for a detailed overview see (57)). Filamentous phages isolated from the Kiel *V. alginolyticus* ecotype share more homology with filamentous phages isolated from *V. parahaemolyticus* than with other non-Kiel *V. alginolyticus* strains. This suggests a constant movement of filamentous phages between different *Vibrio* species without losing the ability to replicate in the old host(s). Indeed, some filamentous vibriophages have a very broad host range (20) and movement of vibriophages is not uncommon (58-60). If filamentous phages are able to establish a chronic infection in the new host, this movement of phages across species boundaries will facilitate horizontal gene transfer (HGT), which plays a significant role in the evolution of vibrios (57).

HGT also contributes substantially to the emergence of pathogenic vibrios from non-pathogenic environmental populations (57). For instance, CTXΦ is able to transduce the cholera toxin (CT) from *V. cholera* to *V. mimicus* leading to the emergence of a pathogenic *V. mimicus* form (58, 59). Many vibriophages contain virulence genes responsible for severe gastro-intestinal diseases (47, 48). For instance, almost 80% of clinical *V. parahaemolyticus* strains contain filamentous phages, encoding the zona occludens toxin (Zot) (22). Also, non-human pathogens, such as *V. coralliilyticus* and *V. anguillarum* contain prophage-like elements encoding Zot, suggesting frequent horizontal gene transfer (HGT) of Zot via prophages among vibrios (11). In the present study, we found one filamentous phage, i.e. *Vibrio* phage VALGΦ6, that contains the virulence cluster comprising Ace and Zot, which are common among vibrios (61). Considering the high homology between *Vibrio* phage VALGΦ6 and phages isolated from *V. parahaemolyticus* this might represent another example where the movement of a filamentous phages across species boundaries leads to the transfer of virulence factors possibly being responsible for the pathogenicity of Kiel *V. alginolyticus* ecotypes. Controlled infection experiments revealed a close link between virulence and coverage of the region encoding for *Vibrio* phage VALGΦ6. Strains, for which we observed a strong reduction in the coverage for the region encoding for *Vibrio* phage VALGΦ6 caused a reduced infection load compared to strains, with a high coverage for this locus. This suggests that the low coverage may result in a reduced number of viral particles and potentially a reduced production of both toxins, which may ultimately result in lower virulence. We are aware, that to be able to ultimately prove that Ace and Zot encoded on *Vibrio* phage VALGΦ6 are causing the virulence of our isolates we would need a strain that does not contain *Vibrio* phage VALGΦ6 for further experiments.

## Conclusion

By characterizing two novel filamentous vibriophages isolated from environmental strains we increase our knowledge about filamentous vibriophages, which is as of October 2019 heavily biased towards human pathogens. We show that also non-human pathogenic vibrios represent a reservoir of filamentous phages, which can contain virulence factors and potentially move between species leading to the emergence of pathogens. We want to encourage future studies on the phage-repertoire and their virulence factors of other non-human pathogenic vibrios. By looking at a wider range of *Vibrio* species we will then considerably expand our knowledge on the types of MGEs in *Vibrio* and in particular how they influence the virulence and evolution of this species.

## Author statements

### Authors and contributors

Methodology: RH, Conceptualisation: CCW, HL. Validation: CCW, HL. Formal analysis: CCW, CMC. Data curation: HL. Writing-Original Draft Preparation: CCW. Writing-Review and Editing: CMC, RH, MH, HL, CCW. Visualisation: MH, CCW. Supervision: HL, CCW. Project administration: CCW. Funding: CCW.

### Ethics approval

Approval for using pipefish during infection experiments was given by the Ministerium fu□r Landwirtschaft, Umwelt und ländliche Räume des Landes Schleswig-Holstein.

### Consent for publication

This work does not need any consent for publication.

## Acknowledgements

We thank Jelena Rajkov, and Olivia Roth for useful comments on a previous version of this manuscript.

## Data bibliography

The accession numbers of the eight *Vibrio algionolyticus* genomes analysed in the present study are provided in Supplementary information, Table S1.

The accession numbers of the two newly discovered filamentous phages are provided in the manusrcipt, see Data statement.

The accession numbers of all other filamentous phage genomes used for comparative genomics in the present study are provided in Supplementary information, Table S3 and Figure 5.

