## Supplementary material for "Closely related *Vibrio alginolyticus* strains encode an identical repertoire of prophages and filamentous phages"

**Table S1**: *Vibrio alginolyticus* genomes sequenced in the present study. Shown are the different organs and pipefish of isolation, the presence of Inoviridae, and the accession number of both chromosomes and if available of extrachromosomal phage replicons

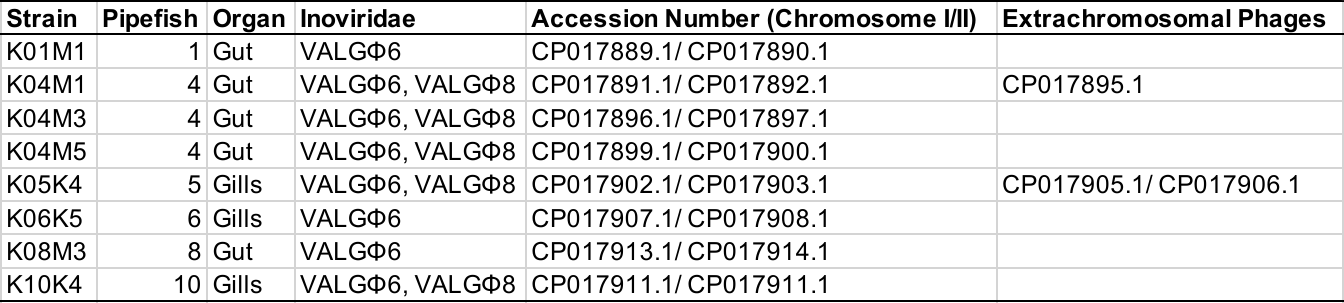

**Table S2**: All prophage regions predicted by PHASTER analysis of each chromosome and the coverage* of Illumina reads generated from phage particles relative to the coverage of the entire chromosome. NA coverage for strain K06K5 is because of missing Illumina sequences for that strain. Chr = Chromosome.

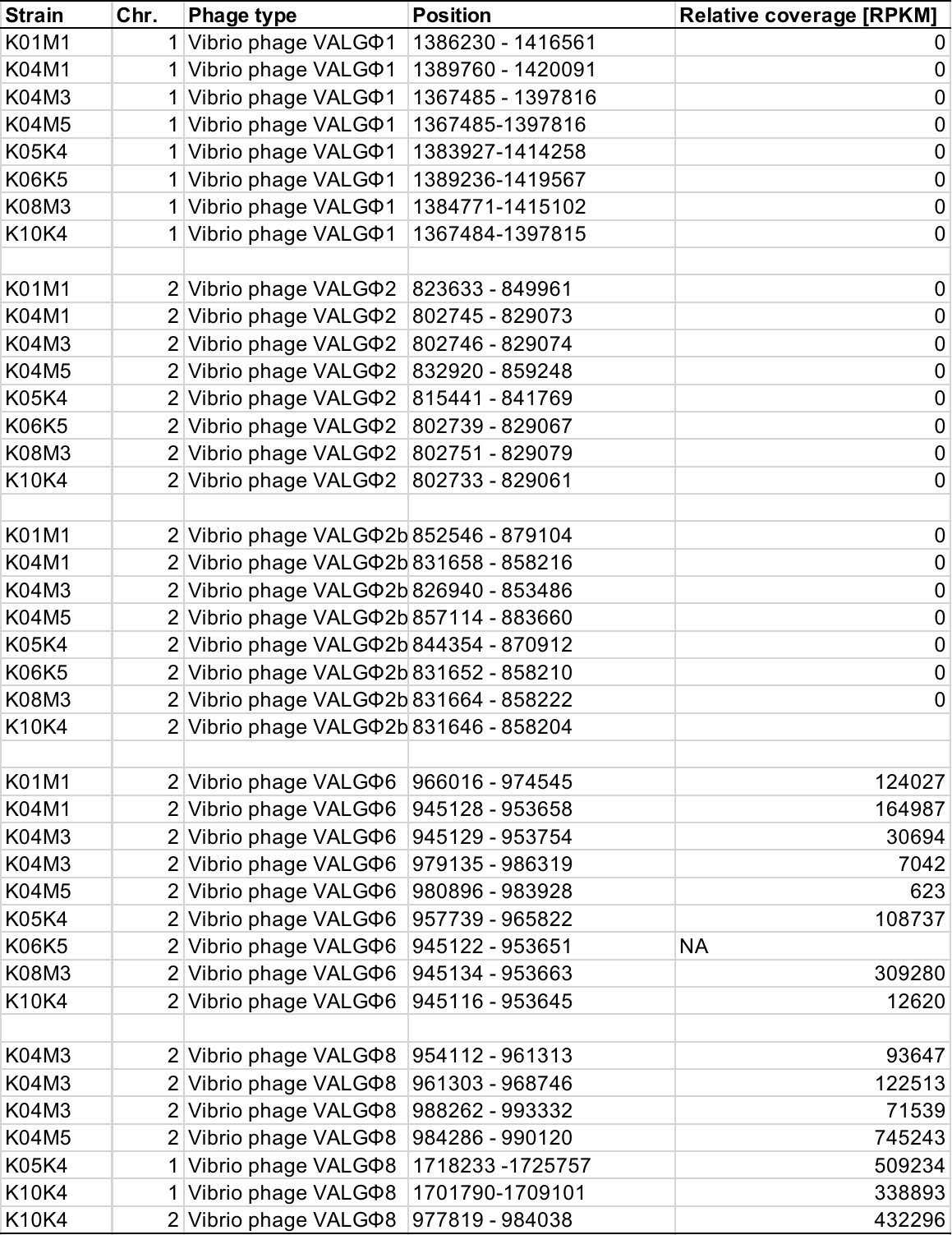

- Coverage values are normalized per region and scaled per million. Unit is RPKM = reads per kilobase million

**Table S3** Filamentous vibriophages used as references for annotation of the Kiel *alginolyticus* phages in the present study

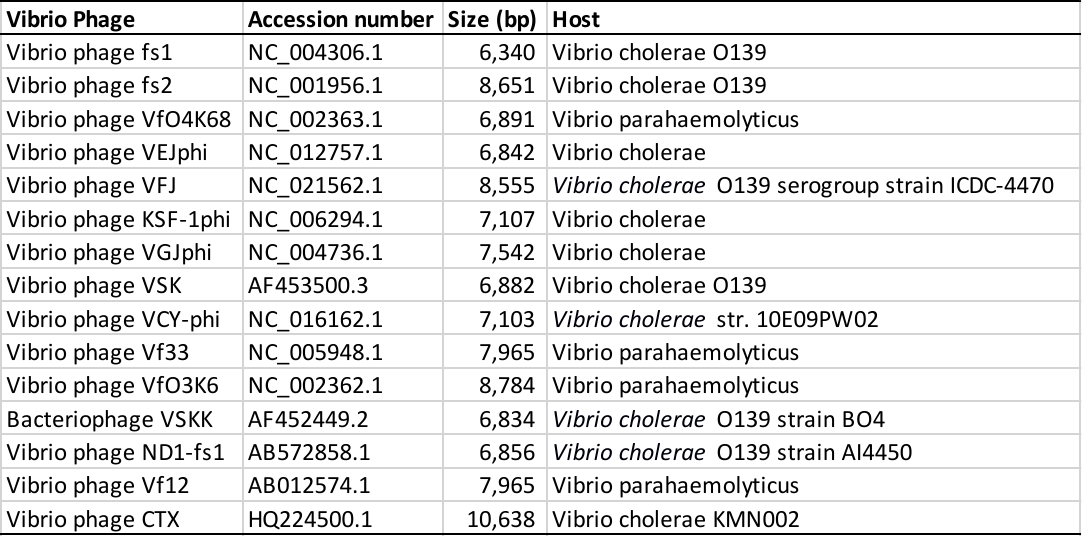

**Table S4**: Prophage regions found in all available closed non-Kiel *alginolyticus* strains predicted by PHASTER and their similarity to Kiel *alginolyticus* phages

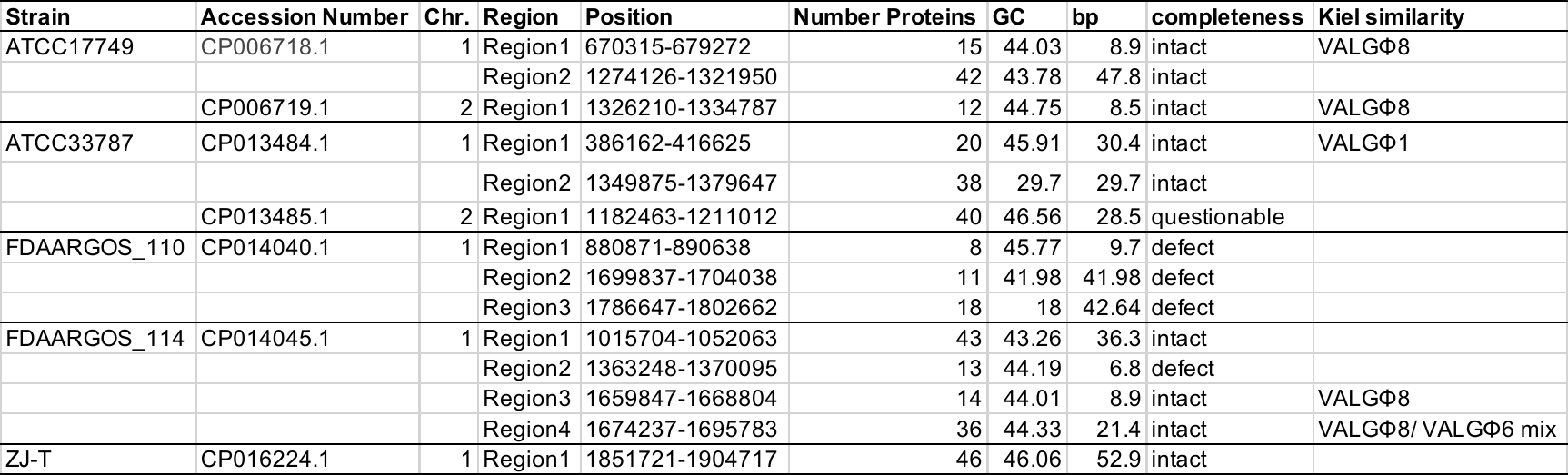

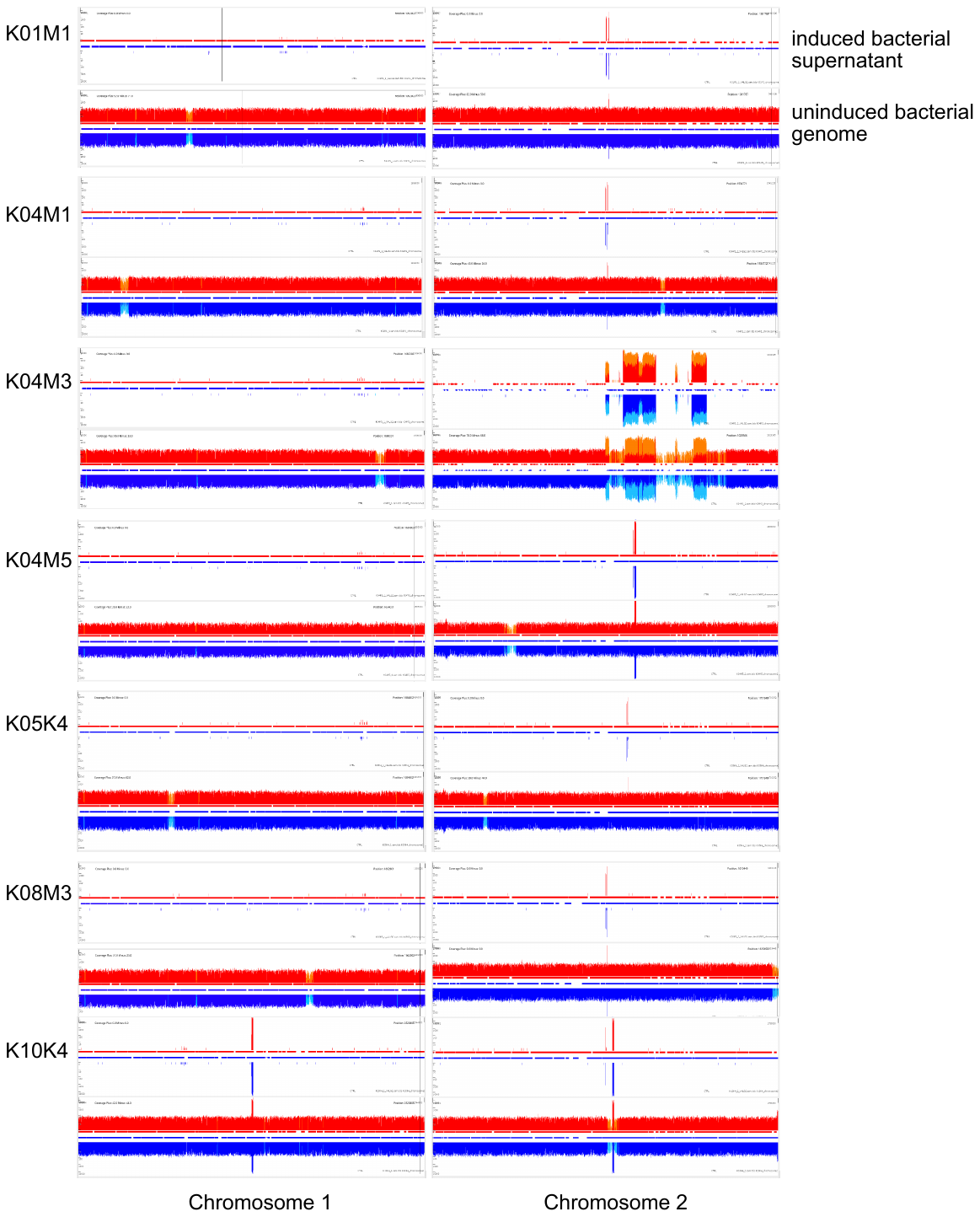

**Figure S1** Coverage (y-axis) for chromosome 1 (left) and chromosome 2 (right) for all eight sequenced strains. Each strain is represented by two images: Top: induced supernatant, bottom: uninduced whole genome sequence of bacteria. Regions of increased coverage correspond to active regions of filamentous phages. Regions of increased coverage in uninduced supernatant are identical with regions of increased coverage in bacterial genome indicating that induced and uninduced cultures produce comparable amounts of filamentous phages.

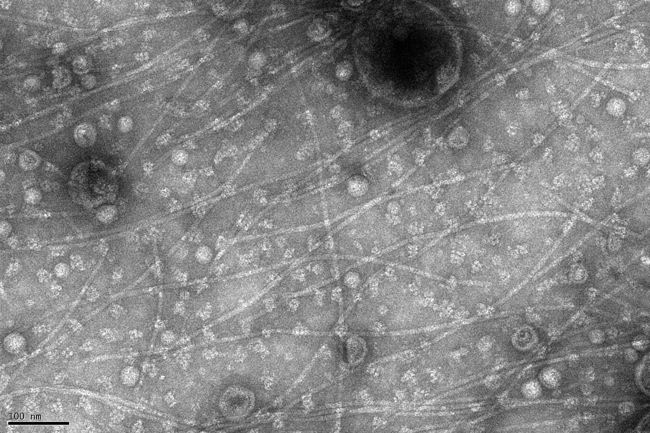

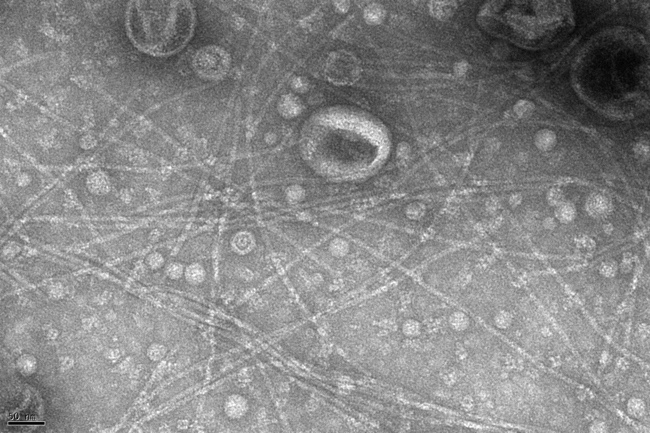

**200 nm**

**200 nm**

**Figure S2** Electron micrographs of filamentous phages of V. alginolyticus K10K4 (Mixture Vibrio phage VALGΦ6 and Vibrio phage VALGΦ8) and V. alginolyticus K01M1 (Vibrio phage VALGΦ6). Arrows point to single filaments.

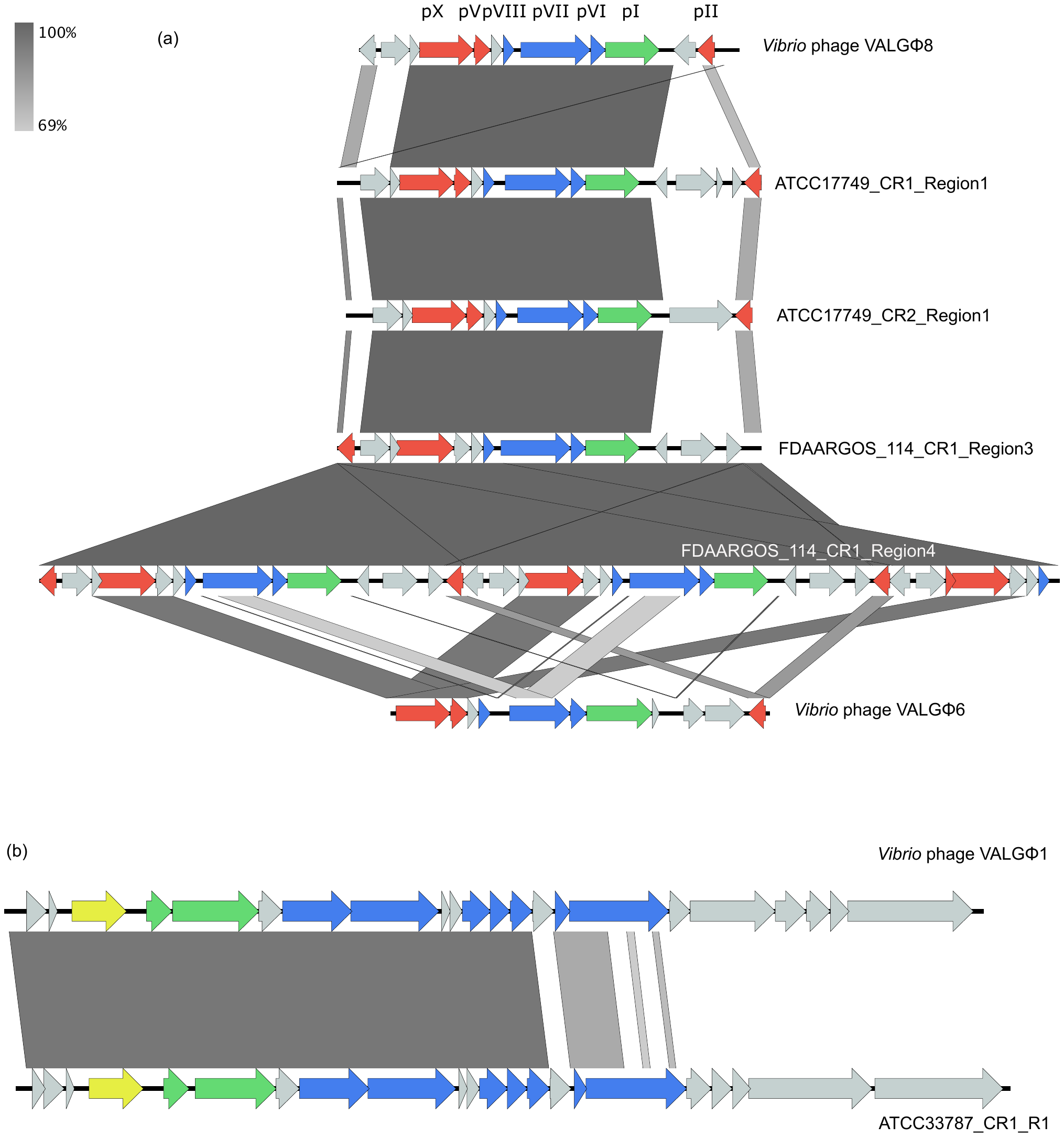

**Figure S3** Whole genome alignment of predicted prophage regions from non-Kiel V. alginolyticus strains and phages identified in the present study. (a) Predicted prophage regions that show similarity with Vibrio phage VALGΦ8 and Vibrio phage VALGΦ6, ORFs are color-coded according to predicted function for Inoviridae: red: replication, green: assembly, blue: structural proteins, grey: hypothetical proteins. pI – pX correspond to known filamentous phage proteins and putative homologues. (b) Region with similarity to Vibrio phage VALGΦ1. ORFs are color-coded according to predicted function: green: assembly, blue: structural proteins, yellow: integration, grey: hypothetical proteins. High homologous sequences are indicated by dark grey and low homologous sequences by light grey.

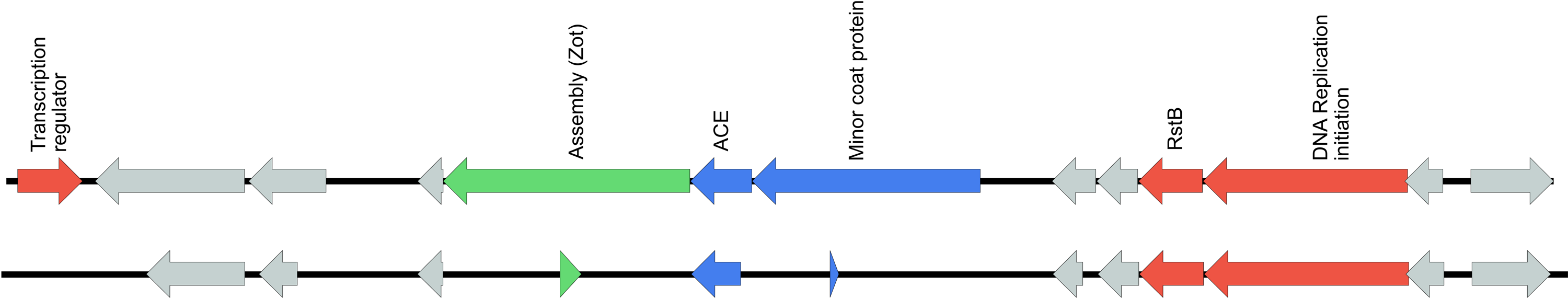

**Figure S4** Genomic maps of both Vibrio phage VALGΦ6 found in strain K04M3, each of which represents the start of a multi-phage cassette. While the first one is complete, the second one has major deletions and no transcription regulator. ORF-coding as in Figure 2.

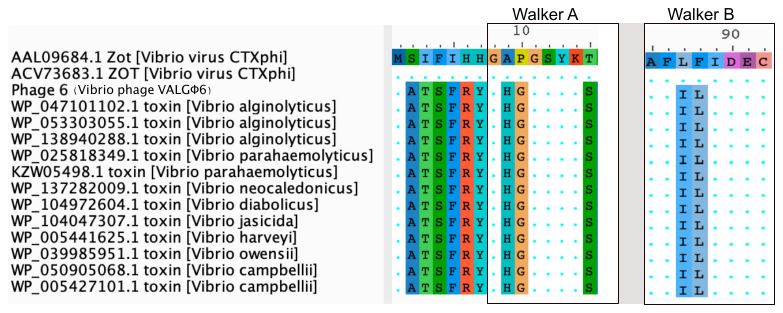

**Figure S5** Zot protein Alignment, regions showing the Walker A and walker B motifs as well as the transmembrane domain on Vibrio phage VALGΦ6 encoded Zot proteins in comparison with other known Vibrio encoded Zot genes. Sequence identity is indicated with dots.

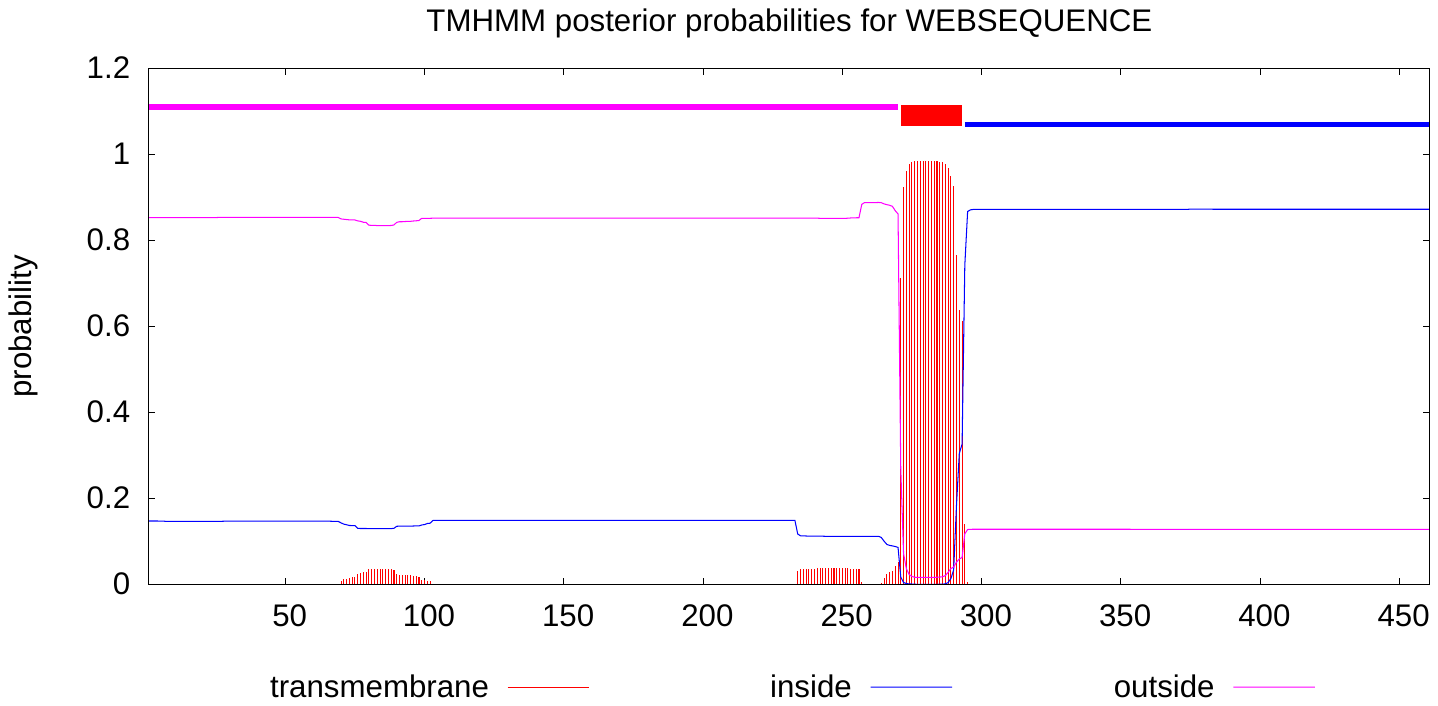

**Figure S6** Prediction of transmembrane domains of the Vibrio phage VALGΦ6 encoded Zot protein.
